# Genetic dissection of tissue composition in genetically diverse mouse populations

**DOI:** 10.1101/2025.10.13.682042

**Authors:** Gregory R. Keele, Matthias B. Moor

## Abstract

**Background:** Single-cell RNAseq data can be harnessed to infer cell-type compositions from spatial or bulk transcriptomes, providing greater biological context underlying systemic tissue or organ dynamics. Here, we used a transcriptome-based systems genetics approach to assess biological factors that influence the cellular composition of tissues, including sex, age and genetic variation in two multiparental mouse resource populations derived from the same 8 parental strains.

**Methods:** We leveraged publicly available bulk RNAseq data from kidney, heart and bone from 188 Diversity Outbred (DO) mice and male/female pairs from 58 inbred strains of the Collaborative Cross (CC). DO mice were aged 6, 12 or 18 months. We used single-cell RNAseq data from healthy C57BL/6 mice as reference for cell-type decomposition of bulk transcriptomes. We tested for differences in cell type composition based on age and sex. We then performed genetic analysis on the cell-type composition phenotypes, estimating the heritability of cell composition levels, mapping cell composition quantitative trait loci (ccQTL), and assessing candidate gene intermediates of ccQTL based on a genetic mediation approach and support in the literature.

**Results:** The heritability of cell-type compositions differed between the inbred CC and outbred DO. Contrasting this, we observed strong and consistent sex differences between the CC and DO. For example, we observed consistency in renal proximal tubule segment 3 (higher in males), T cell abundance in kidney (higher in females), ventricular cardiomyocyte (higher in males) and endocardial cells (higher in females). DO mice exhibited increased levels B and T cells with age, representing an age-related increase in overall renal immune cell content. In heart, DO mice showed decreased ventricular cardiomyocyte composition and increasing endocardial cell content with age. In bone from DO mice, vascular endothelial cells decreased with age and bone marrow stromal cells increased with age. Genome-wide association yielded several cell composition QTLs (ccQTLs) in the three tissues, likely related to known biological phenomena. These include a ccQTL for cardiac fibroblasts in the CC that contains the *Frg2* gene family and fibroblast growth factor inhibitor *Spry2*, and a bone vascular endothelial cell ccQTL in the DO that encompasses 10 genes encoding members of the *S100a* protein family. In addition, our analysis revealed several ccQTLs for different renal tubular segments, immune cell content, and approximately two thirds of cardiac cell types.

**Conclusion:** We establish a role for transcriptome-inferred cellular states or composition for QTL mapping, which adds further utility to existing resources. Furthermore, differences in heritability and QTL between the CC and DO highlight how variation in allele frequencies and inbred vs outbred genetic backgrounds allow for population-specific genetic effects to be discovered. Several of the renal tubular segment and cardiac cell ccQTLs have narrow confidence regions with the potential to harbor novel biological insight. Overall, we propose that our ccQTL mapping approach can be applied systematically across readily available genetic resource transcriptomes to enable new biological discoveries.

## Introduction

With the advent of next-generation sequencing methods for RNA quantification over 15 years ago,(1) high-quality bulk transcriptome profiling is now widely used in both rodent and human experimental systems. In part due to the open-science approach of the computational biology community, a plethora of processing and analysis workflows have emerged for tissue-level data analysis, and also for the more advanced single-cell methods, including single-cell, single-nucleus and spatial transcriptomics. While most methods from bulk transcriptomes have been adopted and modified for single-cell analyses, several more recent single-cell methods have also vice versa been adopted for bulk analyses, including Ligand-Receptor interaction analysis, bulk pseudo-time trajectories, and cell-type deconvolution from bulk samples.(2–4)

The field of genetics has over decades developed statistical methods for harnessing the genomic heterogeneity of natural and experimental populations to identify genomic loci potentially driving or modifying a phenotype, so-called quantitative trait loci (QTL). Like other fields, mouse genetics has leveraged bulk transcriptome data from microarrays or RNAseq to interrogate the effects of both naturally-derived and laboratory-induced genetic variation on genes’ expression from bulk tissue samples in populations such as the BXD, Collaborative Cross (CC) or the Diversity Outbred (DO) population.(5,6) Genetic variants that are associated with gene expression are commonly referred to as eQTLs, which has rapidly grown as a sub-field of genetics that seeks to elucidate the contributions of transcriptional regulation to complex phenotypes.

eQTL mapping has provided tremendous insights into biology, illuminating disease mechanisms and molecular pathways across all kingdoms of life.(7–9) Notably, there is currently are vast resources of transcriptome databases, including many species of plants,(10) mice (GeneNetwork2),(11) rats (https://ratgtex.org/) and humans [GTEx Portal(12), eQTL Catalogue https://www.ebi.ac.uk/eqtl/, and https://seeqtl.org/home/], which are freely and publicly available for further analyses and data mining for new discoveries.

New analytical approaches that leverage many genes from transcriptome data can reveal signal and make inferences beyond the level of single genes and thus potentially hold greater relevance for complex traits. For example, gene co-expression network analysis has been harnessed to define genomic loci associated with gene modules.(13–15) Transcriptome data can be leveraged to address many questions pertinent to complex traits and our understanding of mammalian physiology and disease; examples include: What drives the development or plasticity of different renal tubular segments? What are the biological determinants of bone marrow composition? What are the drivers of heart fibrosis and loss of muscle during aging? The analytical approaches are needed to utilize these rich gene expression resources and advance our understanding of how gene expression regulation contributes to complex traits.

In this study, we address this gap by combining deconvolution transcriptome analyses in genetically heterogenous mice with QTL mapping to identify factors that drive the cell composition of complex tissues. Previous approaches to understand the composition of tissues have used fluorescence-assisted cell sorting (FACS) of blood, followed by QTL mapping of B or T cell lymphocyte subpopulations in human populations(16,17) and mice genetic reference populations.(18) Alternatively, FACS followed by RNAseq has been used to provide cell-specific eQTLs.(19) In a further step, transcriptome-inferred blood cell composition enables eQTLs mapping in specific cell types.(20,21)

In the present work, we extend the prior work done on blood composition to three solid and potentially more complex tissues (kidney, heart, and bone). We demonstrate the pairing of cell composition inference from bulk RNAseq data in multiple organs from two related genetically diverse mouse populations with statistical genetic analyses, producing cell composition heritability, insights into the role of sex and age effects on the inferred cell composition, and a map of ccQTLs that includes known and novel candidate genes associated with phenotypes.

## Methods

### Data and code availability

Unless stated otherwise, we used R version 4.5.1 and RStudio 2025.5.1. Sources of all publicly available datasets are listed. Computational code is available on request to the authors.

### Bulk RNAseq and genotype data retrieval from DO and CC mice

We accessed genotypes and processed bulk RNAseq data from kidney and heart from a previous study(22) representing male/female pairs of 58 CC strains. Gene count data from CC datasets underwent normalization by the TMM method using RNAlysis 3.9.2. In addition, we downloaded the normalized RNAseq data from kidney(23) (GSE121330), heart(24), and bone from 188 DO mice at https://qtlviewer.jax.org/#datasets, along with the metadata age and sex, and genotype data of all individual DO mice. We note that the kidney and heart cohorts largely represent the same mice (except for 3 mice that were filtered from the heart data based on sample quality) whereas only 64 mice with bone data overlap with kidney and heart. Furthermore, the 188 mice with bone data are all female. Variant genotypes from the 8 founder strains of the CC and DO were accessed at https://doi.org/10.6084/m9.figshare.22630642.v1. Bulk RNAseq data had undergone alignment to the respective founder strains’ alleles using the Genotype by RNA-seq (GBRS) software package (https://gbrs.readthedocs.io/en/latest/) to generate optimized gene counts.

### Single-cell RNAseq data analyses as reference for bulk deconvolution

Two single-cell RNAseq datasets were downloaded from Gene Expression Omnibus (GEO), including dataset GSE107585 of murine kidney from 7 healthy C57BL/6J mice (sex-mixed, 4-8 weeks old),(25) and dataset GSE128423 of bone and bone marrow from 8 C57BL/6J (males, 8-10 weeks old).(26) Furthermore, a single-nucleus RNAseq dataset <u>E-MTAB-7869</u> of hearts from three 3-months and three 18-months old male C57BL/6JRj mice(27) was downloaded from the ArrayExpress database of the European Bioinformatics Institute. All data were reanalyzed with R/Seurat version 5.3. We excluded cells with <200 or >2500 feature counts, and cells with >5% mitochondrial RNA content. After standard normalization and unsupervised clustering, cluster identities were annotated based on cluster-specific gene expression and existing literature in kidney(25,28), heart(29,30) and bone[(26) and https://skeletalcellatlas.org].

### Bulk deconvolution by Bisque

All bulk RNAseq data matrices underwent per-sample deconvolution by R/Bisque package 1.0.5 using the reference-based decomposition mode(4) from 10X Genomics reference datasets.

Kidney data were decomposed into 15 cell types using 14846 and 12816 genes in DO and CC datasets, respectively, representing the number of genes detected in both bulk and single-cell expression. We filtered out 370 and 241 zero-variance genes in DO and CC datasets respectively, and 0 non-expressed transcripts in either dataset. The single-cell based reference was established from 4235 cells. Heart data were decomposed into 13 cell types using 16792 and 14841 genes in DO and CC datasets, respectively, representing the number of genes detected in both bulk and single-cell expression. We filtered out 0 zero-variance and 0 non-expressed genes. The single-cell based reference was established from 26650 cells. Bone data were decomposed into 18 cell types using 17005 genes detected in both bulk and single-cell expression. We filtered out 94 zero-variance and 0 non-expressed genes. The single-cell based reference was established from 44100 cells. For simplicity, the results from bulk RNAseq deconvolution analysis are referred to as “cell types” or collectively “cell composition”, though we note that “transcriptome-inferred cell cluster-like cellular state” would be more accurate.

### Heritability estimation

Heritability is the proportion of phenotypic variability in a specific population that is explained by genetics, often quantified based on cumulative SNP variation or pedigree relationships. We estimated heritability for each cell type using the following linear mixed effect model (LMM):

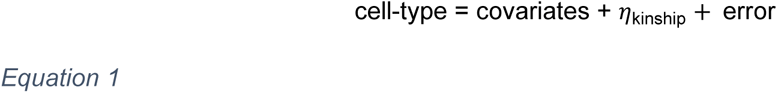

where *η*_kinship_ ∼ N(**0**, **K***τ*^2^) and error ∼ N(0, *σ*^2^) are random terms, representing the effects of cumulative genetic variation and unstructured noise, respectively. **K** is the n-by-n realized genetic relationship matrix (GRM), often referred to as the kinship matrix, estimated from the additive founder allele probabilities at loci spanning the genome.(31) Heritability is then calculated as: 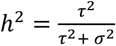. Heritability was estimated using the est_herit function in the R/qtl2 package.(32) Heritability estimates were fit based on the restricted maximum likelihood (REML), which are unbiased for variance parameters.(33)

Cell composition phenotypes were converted to ranks and then transformed into standard normal quantiles (i.e., rank-based inverse normal transformation) to account for non-normal data and reduce the influence of outlying observations. Covariates for the DO kidney and heart data were sex, age as a categorical variable (6, 12, and 18 months), and DO generation as a categorical variable (generations 8-12). Covariates for the DO bone data were the same but with sex excluded. The only covariate for the CC data was sex.

### Testing for age and sex differences

Age and sex differences for the DO datasets were detected using the LMM described in Equation 1. Coefficients for sex and age, now fit as a single continuous variable, were tested using a likelihood ratio test (LRT), based on maximum likelihood (ML) parameter estimates. The fit1 function in the R/qtl2 package was used to fit the LMM.(34) For the CC datasets, we assessed sex differences of cell composition strain-wise using a paired t-test. To declare differences as significantly non-zero, we false discovery rate (FDR)-adjusted the LRT p-values using the Benjamini-Hochberg procedure to account for multiple testing across cell types.(35) A significance threshold of FDR < 0.1 was used.

### QTL analysis

QTL analysis was performed by testing for an association between individual loci spanning the genome and each cell composition trait, R/qtl2’s scan1 function.(34) The underlying model is expanded from Equation 1, adding 8 additive founder allele probabilities at the focal locus as terms that are jointly tested based on ML parameter estimates. The random kinship term in the LMM in Equation 1 was also adjusted to use a leave-one-chromosome-out (LOCO) approach, in which chromosome-specific GRMs are estimated from all markers excluding those on the focal locus’s chromosome, which avoids the random effect and the locus terms from competing and reducing QTL mapping power.(36,37) The strength of association is summarized as a log of odds (LOD) score. We used LOD score > 6 as a lenient threshold, though primarily focus on ccQTLs that met LOD score > 7. We use this approach because converting LOD scores to genome-wide error rates would technically differ between populations and datasets. For reference, in a separate DO cohort of 350 mice, a LOD score threshold of 8.3 controlled the genome-wide family-wise error rate (FWER) at 0.05.(38) A QTL support region was defined using a 1.5 LOD drop.(31)

At detected ccQTL, we characterized the founder haplotype effects as best linear unbiased predictors (BLUPs), using the scan1blups function in R/qtl2, which conservatively constrains unstable coefficients, to identify founder haplotypes driving each QTL and their respective directionality. We also performed SNP association mapping using the scan1snps function in R/qtl2, which converts the eight additive founder allele probabilities to SNP allele probabilities based on the SNP genotypes for the founder strains (https://doi.org/10.6084/m9.figshare.22630642.v1).

### Mediation analysis to identify candidate gene intermediates

For strong ccQTLs with LOD score > 7, we performed a simplified mediation analysis to evaluate genes that are encoded within the ccQTL’s support region and have bulk gene expression data in the data from which cell compositions were derived as candidate mediators. It is important to note that causal interpretation of mediation analysis is generally dependent on strong assumptions, such as the direction of the relationship between the candidate mediator and outcome and whether unobserved confounders may be driving the associations that are being interpreted causally.(39) Generally, these assumptions are not easily evaluated in the context of high throughput experiments of complex molecular systems. Nevertheless, mediation analysis can identify strong candidate intermediates and illuminate the underlying biology driving a detected QTL. Importantly, confirmation of a causal relationship would require experimental follow-up. Given the context of these data, other important caveats include the fact that the true causal gene may not be observed in the bulk gene expression data, or that a bulk-level measurement of its expression does not accurately capture the causal effect driving the cell composition variation.

We used the simplified “LOD drop” approach,(40) which is amenable to omic data in MPP populations like the CC and DO.(41) For each gene with bulk expression data within the ccQTL support region, we fit the peak ccQTL model, as previously described, but now with the candidate gene’s normalized expression data included as a covariate, and the LOD score recorded. Genes that are strong candidate mediators of the ccQTL would presumably represent much of the same statistical signal as the ccQTL, thus their inclusion as a covariate should reduce the ccQTL LOD score. Strong candidate mediators will produce notable drops in the LOD score, likely be encoded near the peak marker for the ccQTL, and possess a bulk tissue *cis* eQTL that co-maps with the ccQTL with similar founder haplotype effects. In addition to mediation analysis, expression of selected candidate genes’ human orthologs was visualized using kidneycellatlas.org/fetal-kidney-developing-nephron.

## Results

Gene expression can be influenced by genetic variation as well as other factors such as sex or aging. In this study, we estimated cell compositions of three tissues and assessed response to these factors in two genetically diverse mouse populations. Eight inbred founder strains, representing both traditional laboratory strains and wild-derived strain, were previously cross-bred to generate the Collaborative Cross (CC) recombinant inbred strain panel and its outbred sister population, the Diversity Outbred (DO) population.(Figure 1A) Here, we performed deconvolution of RNA bulk data into cell compositions of kidney, heart and bone for each individual animal. To this end, we imported, reanalyzed and performed unsupervised clustering of single-cell RNAseq data from the respective tissues of healthy C57BL/6 mice (Figures 1B-D). Clusters were annotated using both cluster-specific differentially expressed genes and well-known cell-specific marker genes from kidney(25,28) (Supplemental Figure 1), heart(29,30) (Supplemental Figure 2) and bone(26) (Supplemental Figure 3).

**Figure 1:**
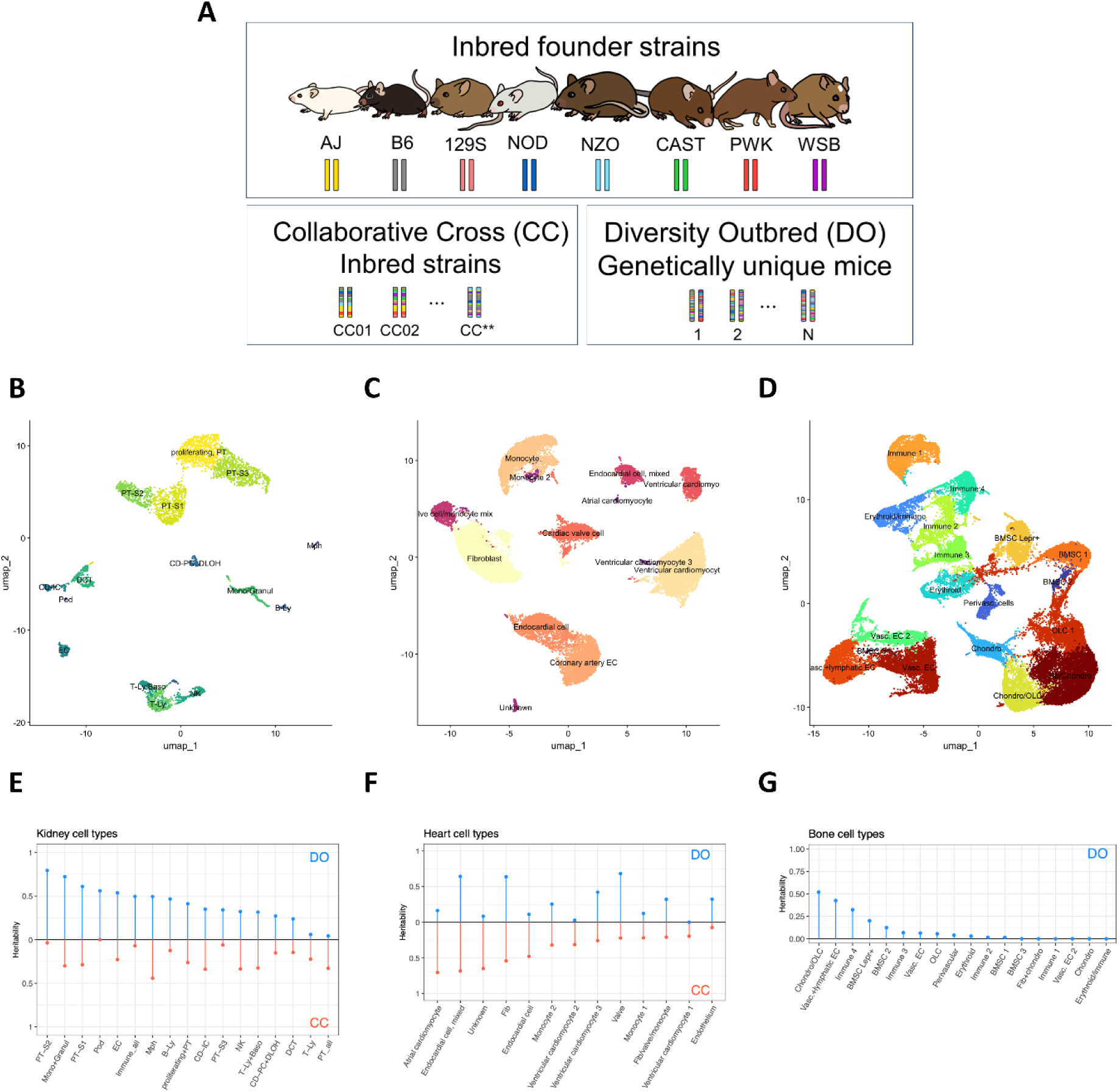
Diagram of the Collaborative Cross (CC) and Diversity Outbred (DO) population (A), which are descended from the same 8 founder strains. The CC are recombinant inbred lines and the DO are highly heterogeneous and each individual is genetically distinct. Cell type clusters from single cell data from mouse kidney (B), heart (C), and bone (D). Heritability for cell composition phenotypes for kidney (E), heart (F), and bone (G). Vertical lines included for clarity. Horizontal line at 0 included for reference.

### Heritability of deconvoluted cell types differs between the CC and DO

After deconvolution of bulk RNAseq data from three tissues using the three single-cell reference datasets, we estimated the heritability (h^2^) of the cell composition traits. In kidney, the DO had notably higher cell composition heritability (mean h^2^ = 41.3%) than CC (mean h^2^ = 21.6%) (Figure 1E). Proximal tubule segment 2 (PT-S2) was the most heritable cell type in DO mice (DO h^2^ = 79.4% vs CC h^2^ = 3.7%) compared to macrophages (Mph) of CC mice (h^2^ = 44.2% vs DO h^2^ = 49.4%). For heart, the CC had the higher average heritability, though the difference was less extreme than in kidney (CC mean h^2^ = 37.4% vs DO mean h^2^ = 29.1%). Valve cells were the most heritable cell type in DO mice (DO h^2^ = 68.2% vs CC h^2^ = 22.1%) and atrial cardiomyocytes in CC mice (CC h^2^ = 70.5% vs DO h^2^ = 16.4%) (Figure 1F). In DO bone (no CC bone data were available), we observed an overall lower heritability profile compared to kidney and heart (mean h^2^ = 10.5%), with the highest heritability found in a mixed chondrocyte / osteoblast (Chondro/OLC) lineage cell cluster (DO h^2^ = 51.9%) (Figure 1G). Overall, heritability for cell compositions from kidney and heart appeared uncorrelated (r = −0.15, p = 0.41) between the CC and DO.

### Sex effects on cell type compositions are conserved between the CC and DO

We next characterized the effects of sex and age on kidney and heart composition (the DO bone cohort only included females). In the kidney, we found substantial sex differences in cell composition that overlapped between the two populations: Proximal tubule segments 2 and 3 (PT-S2 and PT-S3) compositions were more abundant in males, whereas total inferred immune cell content (Immune_all), T-lymphocytes (T-Ly), monocytes and granulocytes (Mono+Granul), a cluster of proliferating proximal tubule cells (proliferating+PT), and podocytes (Pod) were more abundant in females (Figure 2A left panel and Supplemental Figure 4). Overall, sex differences in kidney were significantly correlated between the CC and DO (r = 0.58, p = 0.01). Consistent sex effects in the cell type composition in kidney parallels strong sex differences at the gene level previously observed in kidney tissue from C57BL/6J, (42) CC,(43) and DO.(44)

**Figure 2:**
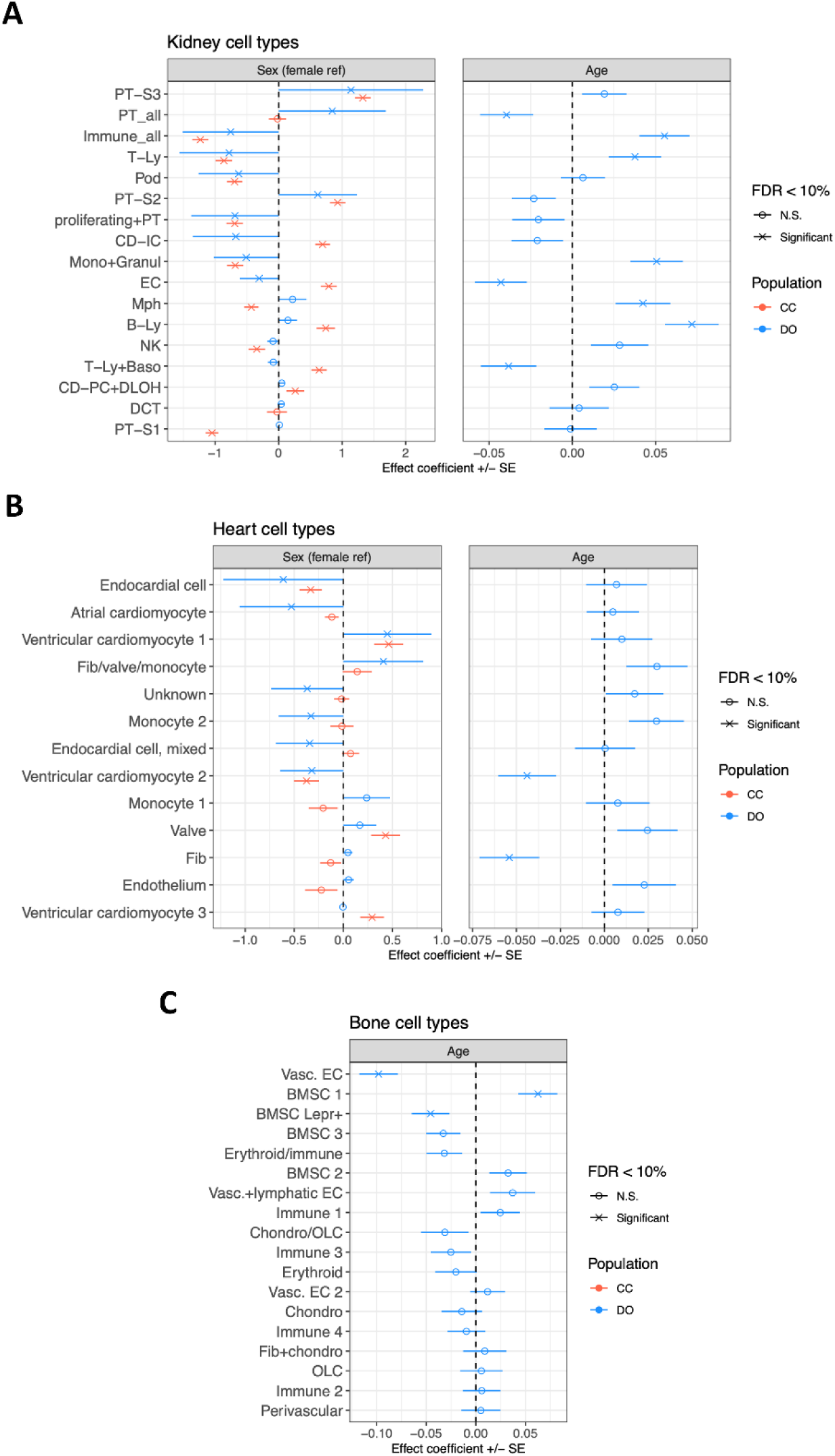
Sex and age differences for cell composition traits from kidney (A), heart (B), and bone (C). FDR_?_ false discovery rate. N.S., not significant.

Similar to kidney, heart cell composition, inferred from transcriptomes, showed some consistent significant sex differences between the CC and DO, with endocardium content (endocardial cell) more abundant in females (Figure 2B left panel and Supplemental Figure 4). An additional non-pure cluster of endocardial cells (endocardial cell, mixed) was more abundant in female DO mice but no sex difference was observed in CC mice. Similarly, atrial cardiomyocytes were more abundant in female DO mice but without a sex difference in CC mice. Ventricular cardiomyocytes showed a more nuanced sex difference with males of both populations harboring higher compositions of “type 1” ventricular cardiomyocyte cluster, whereas females of both populations were more likely to have higher “type 2” ventricular cardiomyocytes compositions. CC males showed higher levels of a valve cell type and ventricular cardiomyocytes of a third cluster, whereas no sex difference was found in the DO (Figure 2B left panel and Supplemental Figure 4). Mirroring kidney, sex differences in heart were correlated between the CC and DO (r = 0.56, p = 0.05).

### Age influences cell type compositions in kidney, heart, and bone

The DO cohorts were from cross-sectional studies of age, consisting of three age groups (6, 12, and 18 months), allowing us to characterize cell composition differences between age groups. In kidney, we detected an increase in immune cell content (Immune_all, T-Ly, Mono+Granul, Mph, and B-Ly) with increasing age (Figure 2A right panel). Cell type compositions in kidney that decreased with age included all proximal tubule cells (PT_all), endothelial cells (EC), and a combined cluster of T-lymphocytes and basophils (T-Ly+Baso). In heart tissue, we observed age-associated declines in cells annotated as fibroblasts (Fib) and a ventricular cardiomyocyte cluster (ventricular cardiomyocyte 2). Several cell types including mixed clusters showed non-significant increase with age (Figure 2B, right panel): a cluster of fibroblast, valve, and monocytes (Fib/valve/monocyte), monocyte (monocyte 2), valve, and endothelium. In DO bone, age effects were detected for a type 1 vascular endothelial cell (Vasc. EC) cluster, which declined with age, and reciprocal age-related differences for bone marrow stem cell clusters (BMSC 1 increase with age and BMSC Lepr+ decrease with age) (Figure 2C). Overall, these data support the utility of cell-type decomposition of tissue-level bulk RNAseq data to detects biological drivers of tissue composition.

### ccQTL mapping analysis reveals population-specific genetic loci

A key aim of this study was map genetic loci that associate with transcriptome-inferred cell composition, which we refer to as cell composition quantitative trait loci (ccQTL). As such, we mapped ccQTL for the whole set of cell composition traits for kidney and heart in both CC and DO and bone in the DO only. Consistent with the general discordance of heritability between the CC and DO, no ccQTL aligned between the CC and DO. For DO kidney, we mapped 21 ccQTLs based on LOD score > 6 and 4 ccQTL based on LOD score > 7 (Figure 3A). These ccQTLs included two strong loci with LOD > 9 for proximal tubule segment 1 (PT-S1) and T lymphocytes (T-Ly) (Figure 3A and Supplemental Figure 5). Additional strong ccQTLs were mapped for segments of the nephron: distal convoluted tubule (DCT) cells and connecting tubule – intercalated cells (CD-IC) (Figure 3A). For CC kidney, we mapped 3 ccQTLs with LOD > 6 (none surpassed LOD score > 7), two for immune cells (T-Ly and Mono+Granul) and one for a mixed cluster annotated as proliferating and PT cells (proliferating+PT) (Figure 3B and Supplemental Figure 5).

**Figure 3:**
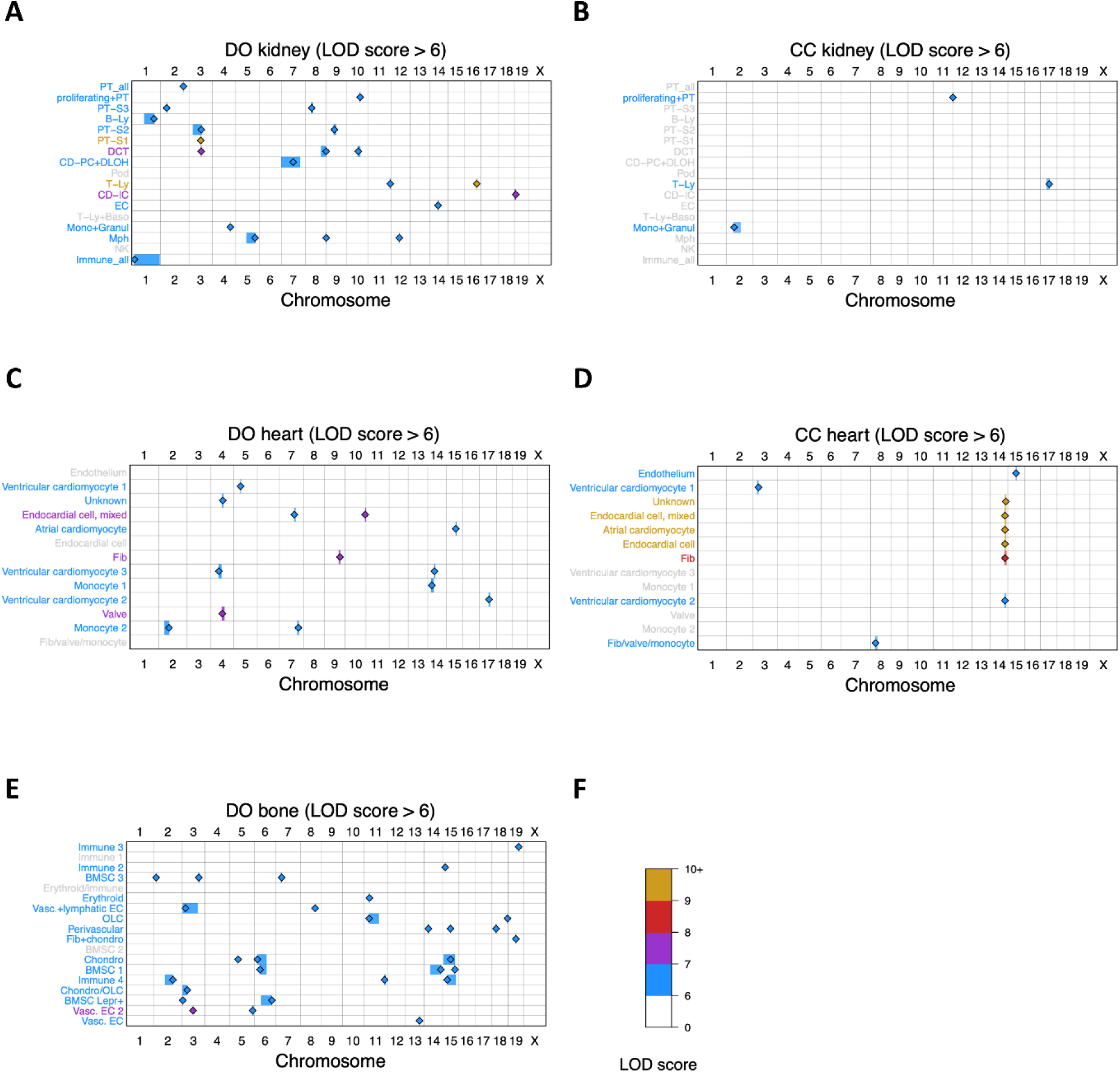
Maps of cell composition quantitative trait loci (ccQ,TL) for DO kidney (A), CC kidney (B), DO heart (C), CC heart (D), and DO bone (E). (F) LOD score color scale. Diamond symbol marks the peak locus of the ccQTL region. Colored shading around diamond symbol represents the QTL support region. Cell composion labels (rows) are colored according to their peak ccQTL.

Next, we assessed ccQTLs in heart tissue from both the CC and DO. In the DO, we detected 13 ccQTLs at LOD score > 6 and 3 ccQTLs at LOD score > 7, representing most of the annotated cardiac cell types (Figure 3C). In the CC, we detected 9 ccQTLs at LOD score > 6 and 5 at LOD score > 7. Notably, those 5 strong ccQTLs (as well as one ccQTL that did not surpass LOD score > 6) all co-mapped to the same region on chromosome 14 and represented two endocardial cell clusters, fibroblasts (Fib), atrial cardiomyocytes, ventricular cardiomyocytes, and an unknown cell-type cluster (Figure 3D).

Finally, for DO bone, we mapped 29 ccQTLs at LOD score > 6 but only 1 ccQTL at LOD score > 7, which was for type 2 vascular endothelium (Vasc. EC 2). We performed a sensitivity analysis in which we combined cell composition traits with related cellular identities (i.e., immune cell clusters 1 to 4, vascular endothelium cell clusters 1 and 2, and all 4 clusters of bone marrow stromal cells), for heart and bone. The resulting ccQTLs did not substantially change (data not shown). To summarize, we detected several high confidence ccQTLs and additional broader, suggestive ccQTLs across kidney, heart and bone from two genetically heterogenous mouse populations. Next, we focus on characterizing the strong ccQTLs and identifying potential candidate genes underlying them.

### Deeper dissection of strong ccQTLs reveals candidate genes

#### PT-S1 ccQTL in DO kidney

The PT-S1 ccQTL mapped to 99.1 Mbp (support interval: 98.1-100.3 Mbp) on chromosome 3 (Figure 4A and Supplemental Figure 6A) and had a peak LOD score of 10.8. Characterizing the founder haplotype effects at the peak locus revealed a high WSB effect, suggesting that DO mice that possess the WSB allele at the locus exhibited higher PT-S1 composition. We performed a mediation analysis, referred to herein as LOD drop(45) (Methods), to assess individual genes as candidate mediators based on their bulk gene expression. Top mediation candidates were *Gm43189*, *Hmgcs2*, and *Wdr3* (Supplemental Figure 7). We also identified *Tbx15*, *Notch2* and *Hmgcs2* based on biological support. i) *Tbx15* was located at the peak LOD score and encodes a transcription factor involved in mesodermal development that promotes glycolytic metabolism over mitochondrial respiration in muscle and pre-adipocytes.(46,47) The human ortholog *TBX15* shows traces of developmental expression in the human nephron and nephron precursor cells (Supplemental Figure 8), including the cap mesenchyme. ii) *Notch2* is a highly relevant developmental gene for PT-S1. The human ortholog *NOTCH2* is expressed throughout the developing human nephron (Supplemental Figure 8), and its variants are responsible for impaired nephron elongation, disrupted ciliogenesis and development of cysts in proximal tubular compartment of human kidney.(48) iii) Hmgcs2 encodes 3-hydroxy-3-methylglutaryl-CoA synthase 2, an enzyme important for mitochondrial function that may preserve renal health.(49) The human ortholog HMGCS2 is expressed in the developing nephron (Supplemental Figure 8).

**Figure 4:**
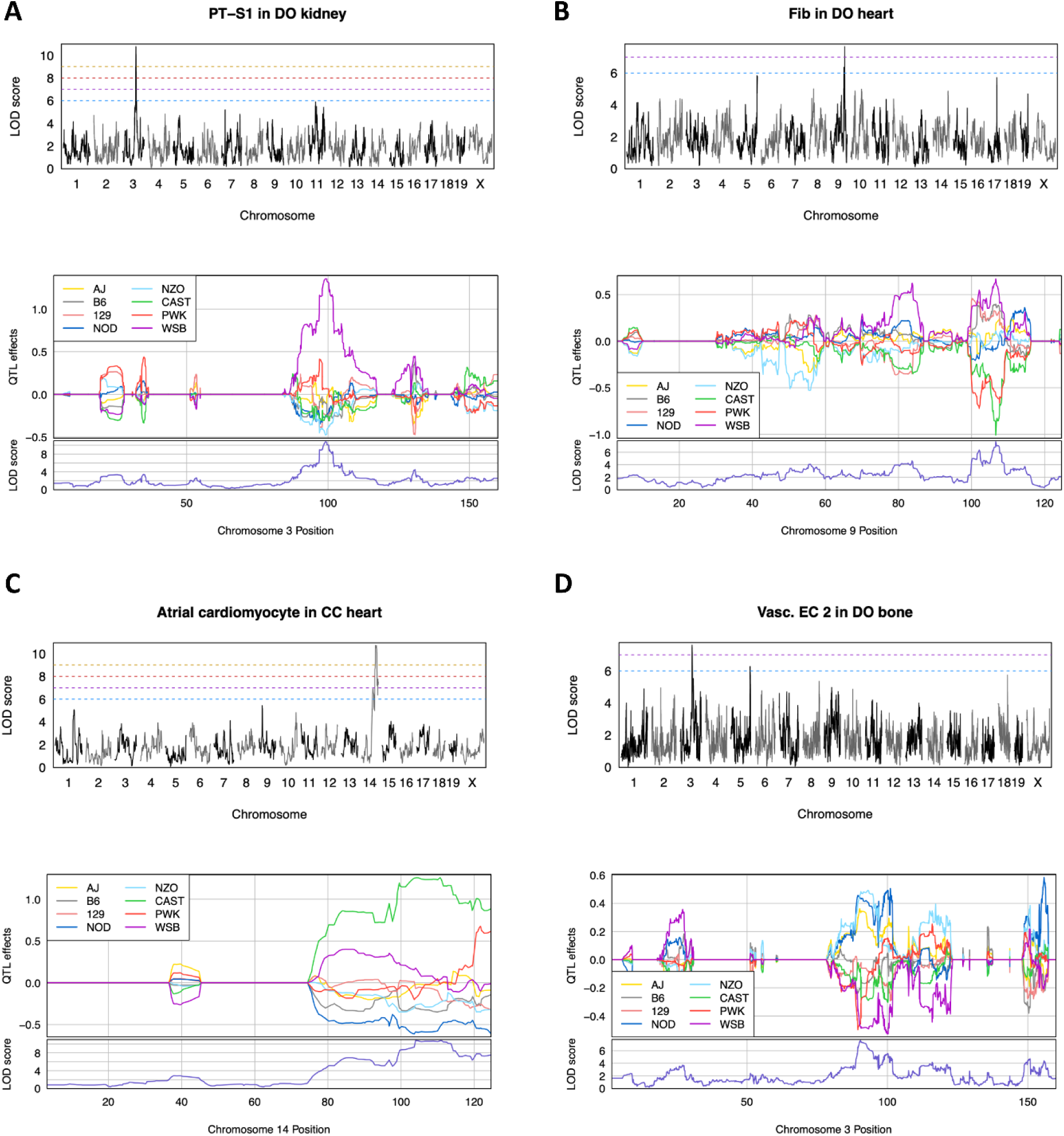
Genome scans (top) and founder haplotype effects (bottom) for PT-SI ccQTL on chromosome 3 in DO kidney (A), Fib ccQTL on chromosome 9 in DO heart (B), atrial cardiomyocyte ccQTL on chromosome 14 in CC heart (C), and Vase. EC 2 ccQTL on chromosome 3 in DO bone (D). Horizontal dashed lines represent LOD score thresholds that each ccQTL surpasses, matching the color scale used in Figure 3.

Additional potential candidates at the PT-S1 ccQTL on chromosome 3 include 6 members of the *Hsd3b* gene family, which encode the 3β-hydroxysteroid dehydrogenases that are crucial for adrenal synthesis of steroid hormones. These include aldosterone, which controls renal salt and water handling.(50) Overall, the high-confidence ccQTL for PT-ST1 covers promising leads for further experimental exploration. Two additional high-confidence ccQTLs for T-Ly and CD-IC in renal tissues of DO mice are described in Supplemental Figure 9A-B.

#### T-Ly ccQTL in CC kidney

The suggestive T-Ly ccQTL mapped to 60.8 Mbp (support interval: 43.1-65.4 Mbp) on chromosome 17 and had a peak LOD score of 6.3 (Supplemental Figure 5C) that broadly covers the major histocompatibility complex (MHC) in mice.(51) Though statistical support for the ccQTL was only suggestive, the MHC represents a critical component of the immune system and thus plausibly regulates the composition of T-lymphocytes in kidney tissue. Furthermore, the MHC possesses high levels of genetic variability, which is potentially reflected in the multi-allelic effect pattern observed for the T-Ly ccQTL (Supplemental Figure 9C).

#### Fib ccQTL in DO heart

The fibroblast ccQTL mapped to 106.6 Mbp (support interval: 101.9-107.7 Mbp) on chromosome 9 (Figure 4B and Supplemental Figure 6B) and had a peak LOD score of 7.7. The haplotype effects at the ccQTL were characterized by low effects for CAST and PWK, intermediate effects for AJ, NOD, and NZO, and high effects for B6, 129, and WSB (Figure 4B). The strongest LOD drop gene was *Nphp3*. The ccQTL region is gene rich and will require follow up to prioritize gene candidates.

#### Co-mapping ccQTLs in CC heart

We detected a genomic region on chromosome 14 (peak locus at 104.0 Mbp with support interval: 103.1-114.2 Mbp) that regulated 6 cell clusters (atrial cardiomyocyte; endocardial cell; endocardial cell, mixed; Fib; unknown; and ventricular cardiomyocyte 2) (Figure 3D). In terms of the atrial cardiomyocyte ccQTL, The CAST haplotype was associated with high compositions (Figure 4C). Atrial cardiomyocyte composition was positively correlated with the other cell compositions, except for fibroblast, which as expected, had inverted haplotype effects at the ccQTL, i.e., CAST haplotype was associated with low fibroblast composition.

LOD drop analysis identified *Gm10076* as the strongest candidate mediator in the bulk expression data (Supplementary Figure 7C). As expected, *Gm10076* co-mapped a strong *cis* eQTL to the ccQTL and exhibited highly consistent haplotype effects (Supplementary Figure 7E). *Gm10076* has recently been investigated deeply and associated with the ribosomal intersubunit bridge protein RPL41(52). Its perturbation was found to have strongly impacted on translation and cell proliferation. Other intriguing genes in the ccQTL region included *Spry2*, a well-described negative regulator of fibroblast growth factor receptor signaling, and the *Frg2* gene family.

#### Vasc. EC 2 ccQTL in DO bone

The vascular endothelial cell ccQTL mapped to 90.4 Mbp (support interval: 89.4-93.6 Mbp) on chromosome 3 (Figure 4D and Supplemental Figure 6D) and had a peak LOD score of 7.6. The haplotype effects largely reflected a bi-allelic pattern with a high group composed of AJ, NOD, and NZO haplotypes and a low group composed of B6, 129, CAST, PWK, and WSB (Figure 4D). Among the genes in the ccQTL region (Supplemental Figure 6D), 12 are members of the *S100a* gene family. Four of the five top LOD drop candidate genes are *S100a* genes (*S100a3*, *S100a4*, *S100a9*, and *S100a14*) (Supplemental Figure 7D). Some *S100a* genes, such as *S100a8*, *S100a9* and *S100a12*, have an established role in stress responses by endothelial cells.(53,54) Notably, the peak LOD score is highly proximal to the *S100a* genes, both in terms of haplotype association, (Figure 4D), SNP association (Supplemental Figure 6D), and the LOD drop analysis (Supplemental Figure 7D), providing strong support for these genes as candidates contributing to the phenotypic variation in vascular endothelial cell composition.

## Discussion

In this study, we used cell-type deconvolution from bulk RNAseq data to determine the relative fraction of cellular states that we consider as a proxy for cell types. Using the obtained cell-type data, we characterized sex effects in two related mouse populations that are largely congruent and consistent with established literature on sex-biased PT-S3 segment cell transcriptomes in the kidney. Further, we establish a framework for cell composition QTL (ccQTL) mapping in complex mammalian organs and identified several ccQTLs, including for renal cell types PT-S1 cells and T lymphocytes, A sex bias of the immune cell landscape in the kidney has previously been observed in different contexts, including in human patients.(55) Overall a sex bias in innate and adaptive immune system is well-established.(56) In addition, age-dependent increases in immune cell proteins or transcripts have previously been described in the kidney,(23,57) corroborating our findings. Furthermore, we describe a sex difference in PT-S3 transcripts. Several markers of PT-S3 are sex-biased in mice,(28) hence the annotation may explain parts of the observed sex difference in PT-S3 abundance in CC and DO mice.

For sex differences in heart tissue, the higher composition of endocardium in from both populations was notable because, to our knowledge, this finding has not previously been reported. It may relate to the finding that approximately 14-25% of endothelial transcriptome is sex-biased(58) and with an expected larger cardiomyocyte mass in males, a reciprocal larger fraction of endocardium may be thus found in females.

The ccQTLs from this study can provide hypotheses that can be explored in future studies. For example, the connection between the WSB-specific genetic variants at the strong ccQTL for PT-S1 warrants further focus. Steroid hormone aldosterone is synthesized with the assistance of proteins from the *Hsd3b* gene family, and aldosterone controls salt and water handling by the kidney. Thus, variants in *Hsd3b* genes may potentially cause compensatory effects on renal salt handling by the proximal tubule. Indeed, in a survey of 40 mouse strains, including 7 of 8 CC/DO founder strains, the WSB strain was among those preferring to consume the least amount of sodium.(59) These promising leads are a good starting point to be addressed experimentally.

Another example of promising finding for future investigation is the Vasc. EC 2 ccQTL in DO bone. Here the *S100a* gene family was highlighted as the potential genetic driver of bone vascular endothelia abundance and warrants further investigation. In a recent study of bone mesenchymal stromal cells isolated from DO mice and subjected to osteogenic differentiation, genetic variation at the *S100a1* locus was identified as a candidate gene for regulating the transition from osteoblast progenitor cells to mature osteoblasts.(15) As bone formation is in part derived from vascular origin,(60) the potential inverse connection between vascular endothelia and osteogenic differentiation associated with this ccQTL should be further investigated.

For the strong co-mapping ccQTL cluster on chromosome 14 for CC heart, *Gm10076* is an intriguing candidate, identified through our mediation analysis. Causal interpretations of mediation analysis are dependent on many assumptions, including that there are no unknown confounders driving the observed associations, which generally cannot be known or confirmed in the context of a high throughput omics study.(39) Nevertheless, mediation analysis can powerfully identify overlapping signal, which is clearly the case for *Gm10076* and its *cis* eQTL that strongly mirrors the ccQTL. This could stem from linkage disequilibrium (LD) between *Gm10076* and the true causal gene in the ccQTL region. Combined with recent evidence for *Gm10076’s* role in translation and cell proliferation, this finding supports further investigation of this ccQTL and *Gm10076.*(61)

Although sex differences were well conserved between the CC and DO, genetics in terms of heritability and ccQTLs appeared to be highly population-specific. Although both populations generally possess the same genetic variation, genetic drift and selection differ due to homozygous lethal alleles in the CC and meiotic drive in the DO.(62,63) Thus, differences in allele frequencies can occur between the CC and DO. In a previous comparison of genetic effects on protein abundance in the CC and DO, we found strong replication of simple genetic effects, i.e., *cis* protein-QTLs, but more complex effects could be population-specific.(64) Specifically, aggregate effects on protein complexes were detectable in the CC, likely representing non-additive effects like recessivity and epistatic effects. Furthermore, for studies that include CC strain replicates, heritability estimates can absorb these non-additive effects.(65) We also note that these data represent multiple studies conducted by multiple labs at different times, also potentially contributing to differences between populations.

General strengths of this study include the combination of both inbred and outbred populations for two of the assessed tissues, allowing direct comparison of sex and genetic effects across populations. As mentioned above, the inclusion of both inbred and outbred populations allows us to observe the effects of alleles in very different background genetic contexts; for example, rare recessive alleles in inbred strains and robust genetic effects in background of the wide-scale heterozygosity of an outbred population. Further, similar RNAseq datasets are widely publicly available, including several more in these specific genetic reference populations, allowing for further refinement and validation with additional cohorts.

This work also has several limitations to note. Although we find these analyses and results promising, validation of findings through experimental follow-up, using methods such as standard histology, immunohistochemistry, or single-cell RNAseq of tissues from the CC or DO population would be needed to confirm our ccQTL findings. We have previously validated perturbation-induced variation in transcriptome-inferred cellular fractions using automated immunofluorescence quantification on immune cells in mouse kidney samples or by standardized histopathology reports of human kidney biopsies.(66) Others have previously combined a similar deconvolution approach with direct histological validation,(67) which is not feasible using the current transcriptome re-analysis approach. Thus, the limitation remains that the inferred cell types inferred may only represent true cellular identities for the easily discriminated renal tubular segments. For other tissues and cell-types, such as in the bone dataset where mixed cell clusters remained, the inferred “cell types” may rather resemble cellular states, e.g., of BMSC, or of the osteoblast lineages. Future research and validation using independent datasets, populations and computational deconvolution approaches may be required to assess this potential limitation.

In addition to the potential that the cell types represent cellular states rather than true cellular identities, we have only used one of the dozens of competing deconvolution protocols. Bisque has been validated as one of the best methods for deconvolution,(4) but many more approaches are available and new methods continue to be developed. Another limitation is that the sample size of CC mice is relatively low, both in terms of the number of strains and the use of a single animal of each sex per strain. The heart reference dataset is single-nucleus RNAseq, whereas cardiomyocytes have more than 1 nucleus. This could potentially drive false positives in which what appear to be different clusters of cardiomyocytes could actually be differing nuclear states within the same cell. However, the ccQTL mapping results were robust to collapsing clusters of same cell type in a sensitivity analysis. Finally, the reference single-cell datasets of the bone and heart contained only males, hence a replication of findings with sex-mixed reference datasets would be ideal.

## Conclusion

We establish a ccQTL mapping framework for bulk transcriptome-inferred data of different tissues based on single-cell reference data. This work represents a promising use for readily available resource data, and future investigations can refine its approach for more deeply harnessing existing transcriptome datasets from population genetics resources.

## Author contributions

This study was separately conceived by both authors at different times and places, followed by a serendipitous collaboration.

## Acknowledgements

The authors acknowledge the contributors of the data analyzed.

## Data availability

All datasets are available as per accession numbers in the methods section. No new datasets were generated. Raw data are available on request to the corresponding author.

---

**Supplemental Figure 1.**
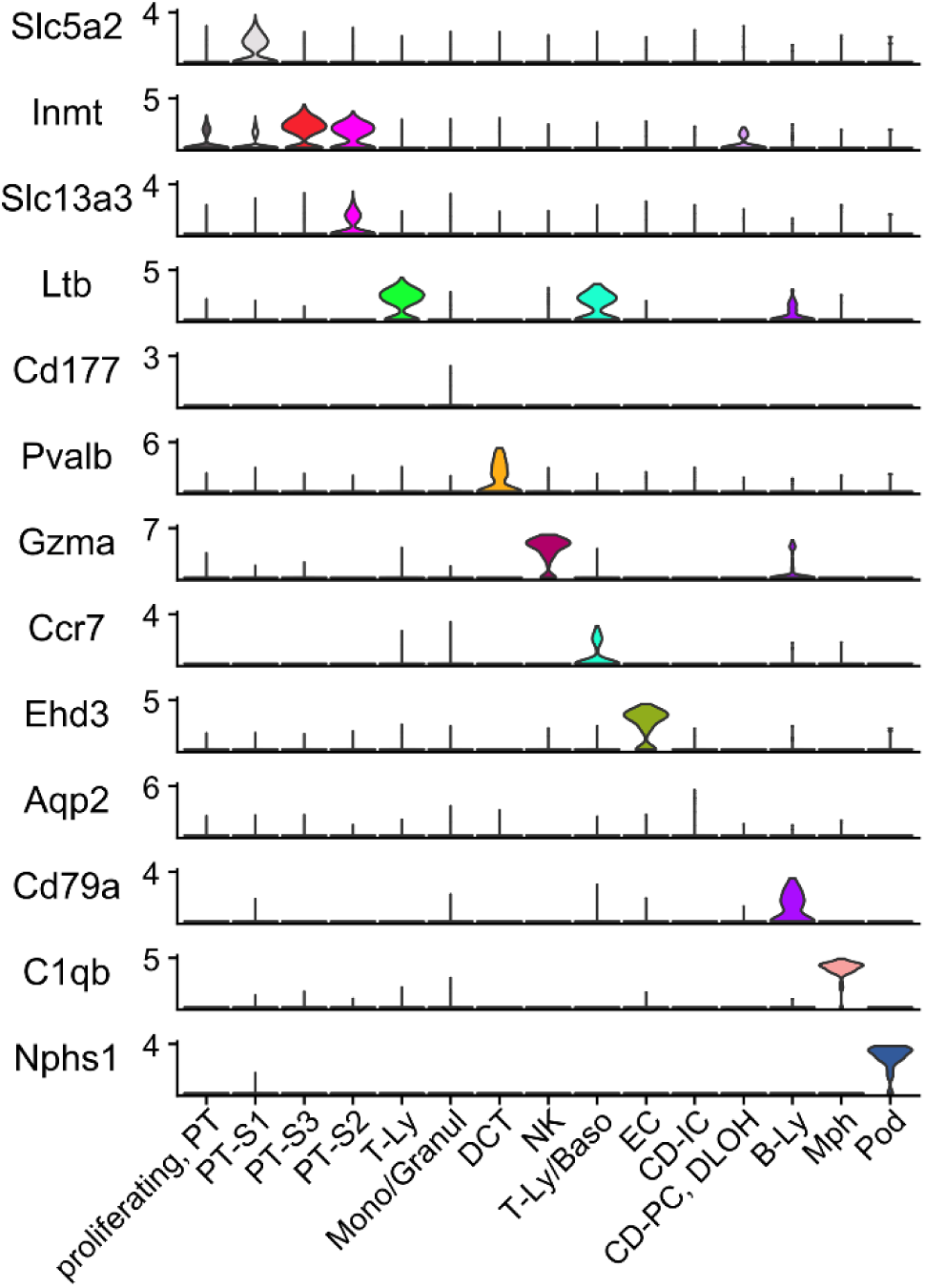
Transcriptional markers for each of the manually annotated clusters of kidney cells from 7 healthy sex-mixed C57BL/6 mice of single-cell RNAseq dataset GSE107585. Ly, lymphocytes. NK, natural killer cells. Base, basophils. Mph, macrophages. Pod, podocytes. EC, endothelial cells. CD-IC, collecting duct-intercalated cells. DCT, distal convoluted tubule. PC, principal cells. DLOH, descending loop of Henle. PT, proximal tubule. S, segment.

**Supplemental Figure 2.**
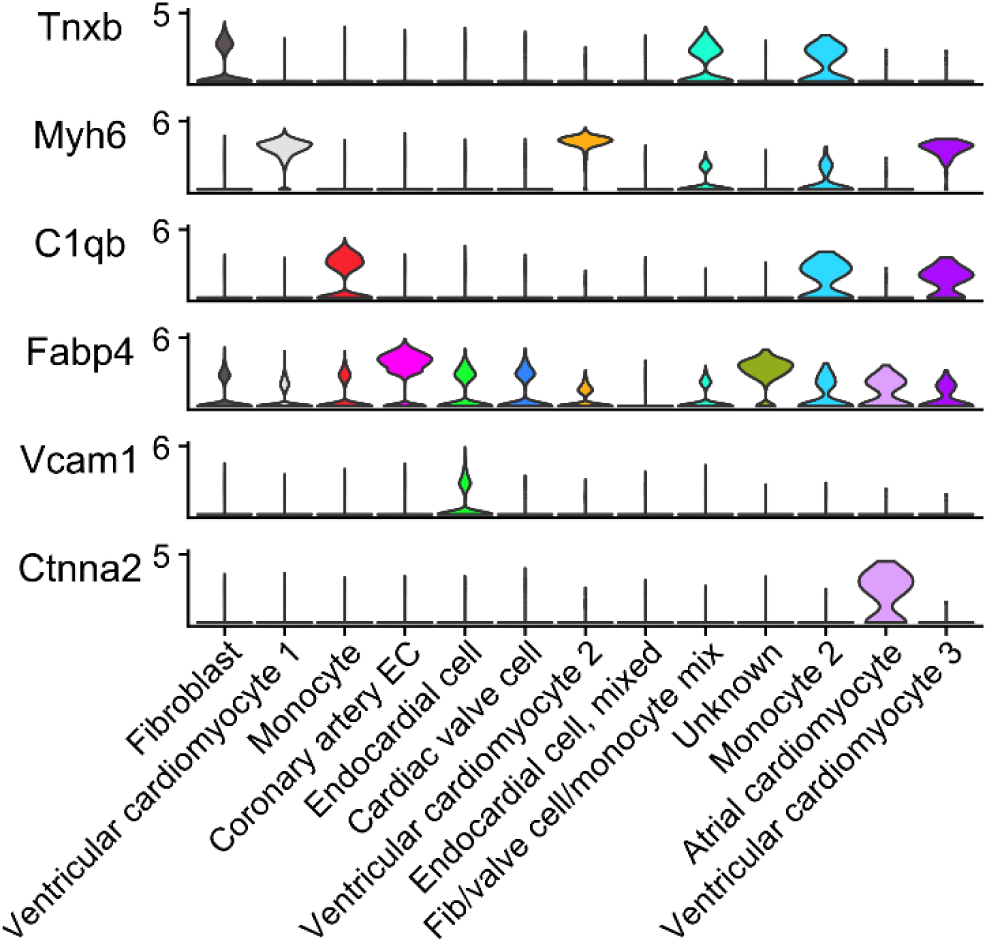
Transcriptional markers for each of the manually annotated clusters of heart cells from 6 male C57BL/6RJ mice of single-nucleus RNAseq dataset E-MTAB-7869. EC, endothelial cell. Fib, fibroblast.

**Supplemental Figure 3.**
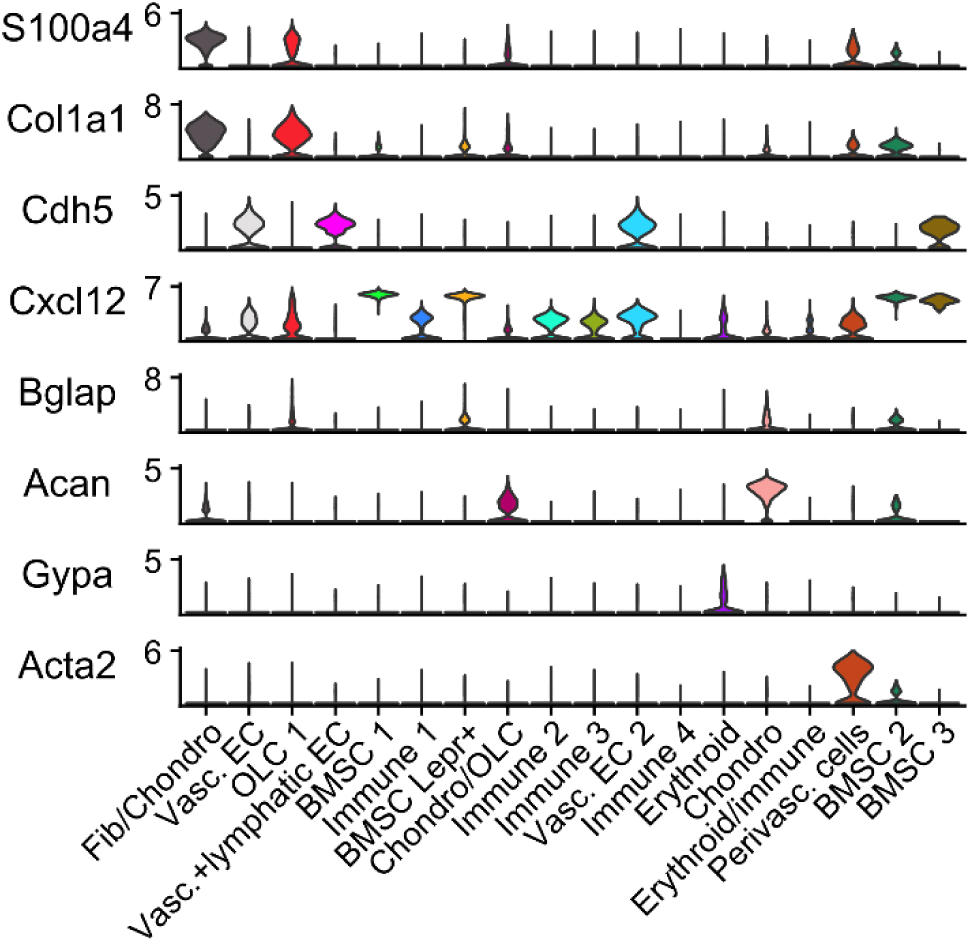
Transcriptional markers for each of the manually annotated clusters of bone and bone marrow cells from 8 male C57BL/6J mice of single-cell RNAseq dataset GSE128423. Fib, fibroblast. Chondro, chondrocyte. Vase. EC, vascular endothelial cell. BMSC, bone marrow stromal cell.

**Supplemental Figure 4.**
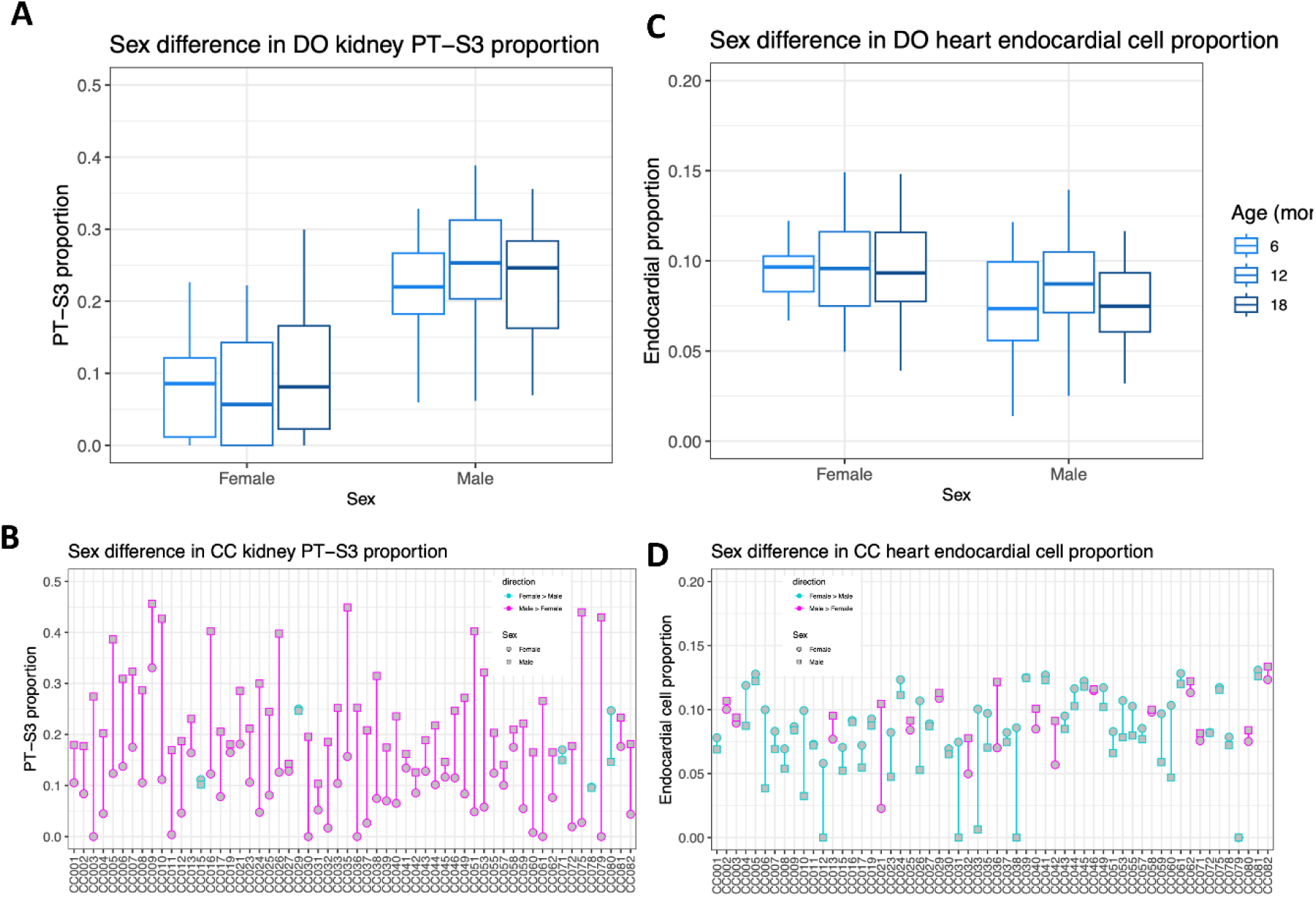
Consistent sex differences, characterized by higher levels in males, observed for PT-S3 in kidney for DO (A) and CC (B). Consistent sex differences, characterized by higher levels in females, observed for endocardial cells in heart for DO (C) and CC (D).

**Supplemental Figure 5.**
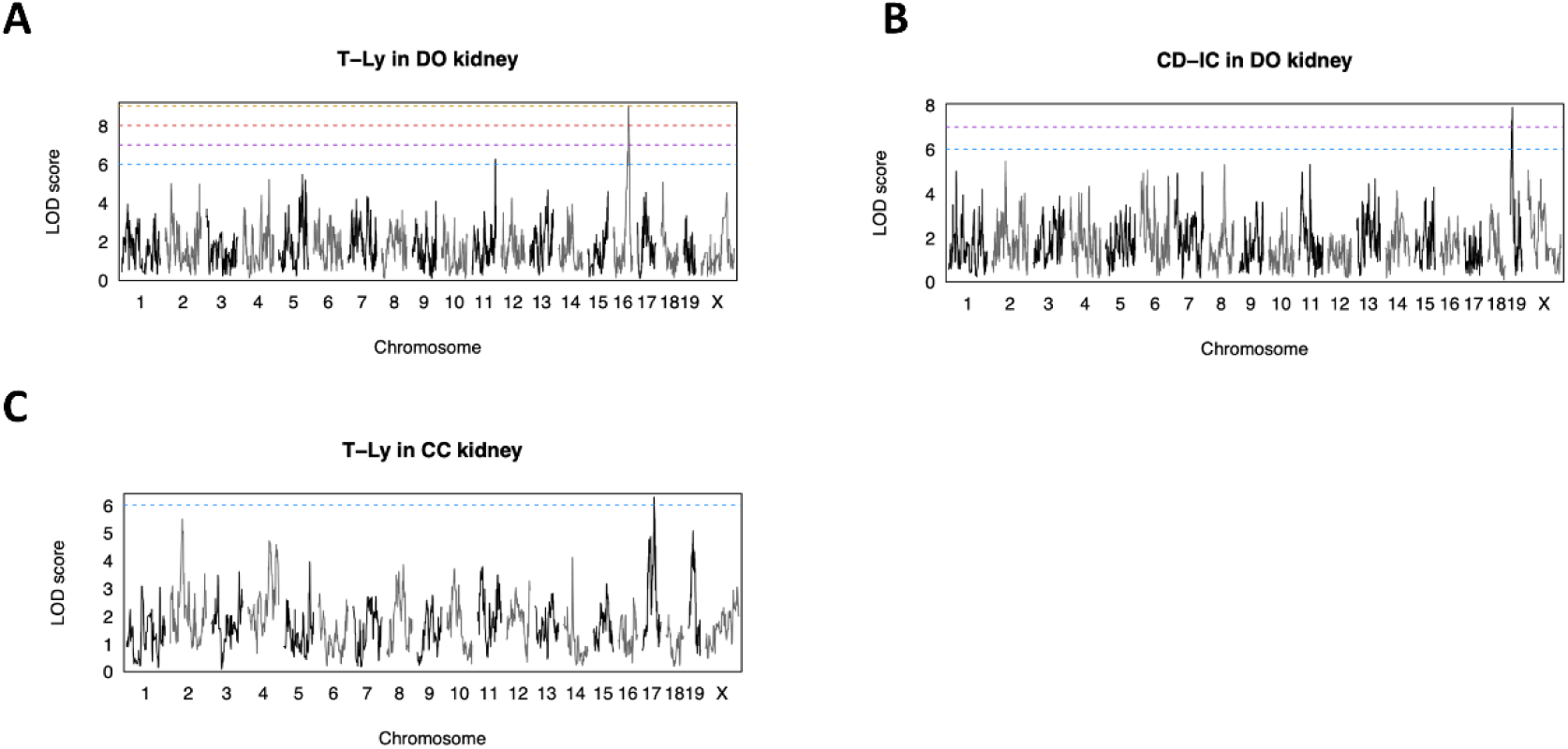
Genome scans for strong T-Ly ccQTL on chromosome 16 (A) and strong CD-IC ccQTL on chromosome 19 (B), both in DO kidney. (C) Genome scan for suggestive T-Ly ccQTL on chromosome 17 QTL that includes the major histone compatability complex locus. Horizontal dashed lines represent LOD score thresholds that each ccQTL surpasses, matching the color scale used in Figure 3.

**Supplemental Figure 6.**
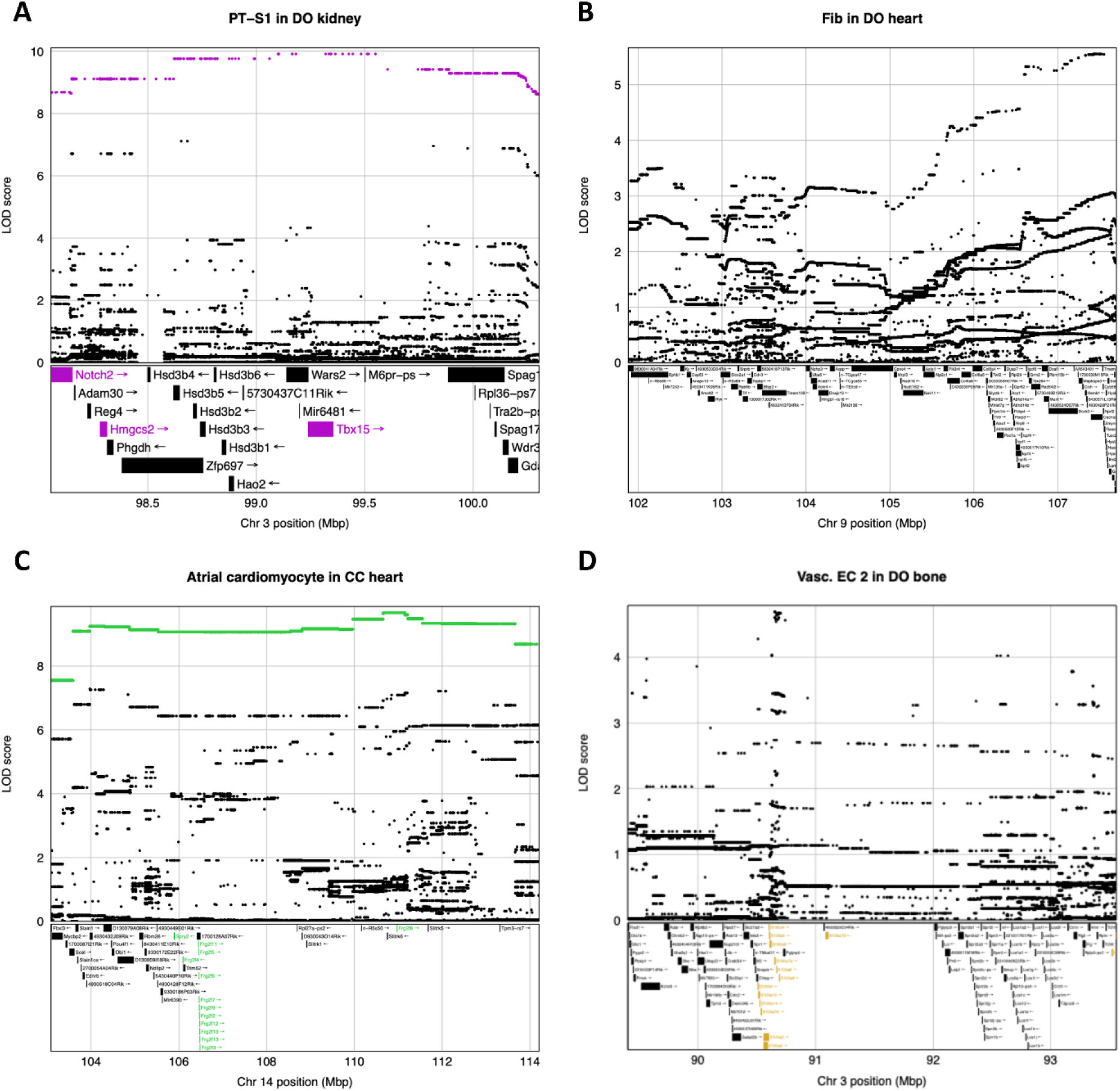
SNP associations and encoded genes in the support regions of ccQTL for PT-SI in DO kidney (A), Fib in DO heart (B), atrial cardiomyocyte in CC heart (C), and Vase. EC 2 in DO bone (D), all highlighted in Figure 3. For the PT-SI ccQTL (Figure 3A), a high WSB haplotype effect was observed (purple). SNPs with a WSB-specific allele and candidate genes are highlighted as purple. For the atrial cardiomyocyte ccQTL (Figure 3C), a high CAST haplotype effect was observed (green). SNPs with a CAST-specific allele and candidate genes are highlighted as green. For the Vase. EC 2 ccQTL (Figure 3D), candidate genes are highlighted as gold. Gene models (gene symbols starting in Gm) were excluded for clarity.

**Supplemental Figure 7.**
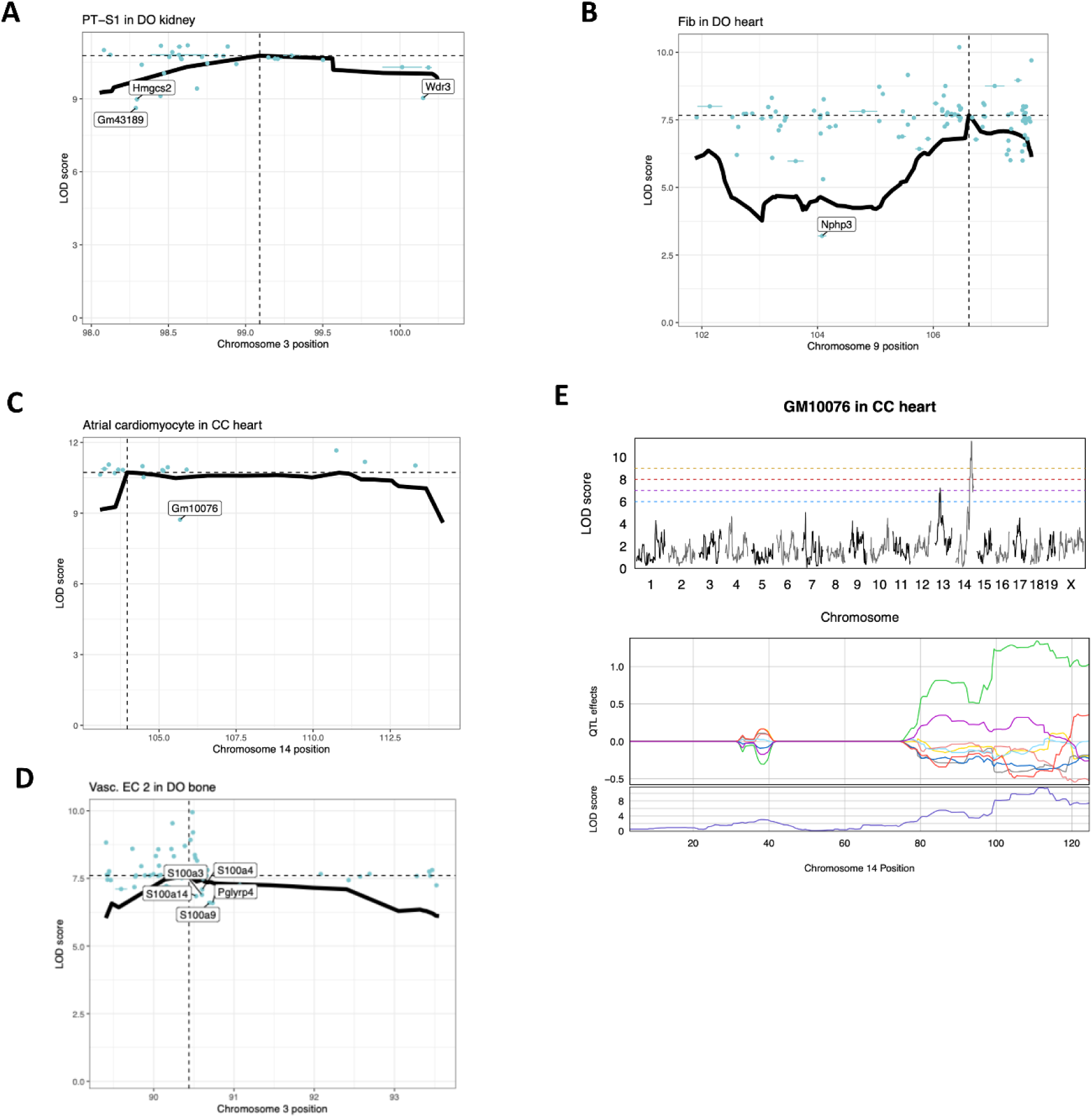
LOD drop results of ccQTL for PT-SI in DO kidney (A), Fib in DO heart (B), atrial cardiomyocyte in CC heart (C), and Vase. EC 2 in DO bone (D), all highlighted in Figure 3. Thick black line represents the LOD score for the ccQTL support region, peak locus marked by the vertical dashed line and peak LOD score by the horizontal dashed line. Candidate mediator genes represented by blue dot and bar. Candidate mediator genes with notable LOD drops highlighted with labels. (E) Genome scans (top) and founder haplotype effects (bottom) for the *cis* eQTL of GmlOO76, a candidate mediator for the atrial cardiomyocyte ccQTL in CC heart. Horizontal dashed lines represent LOD score thresholds that each ccQTL surpasses, matching the color scale used in Figure 3.

**Supplemental Figure 8.**
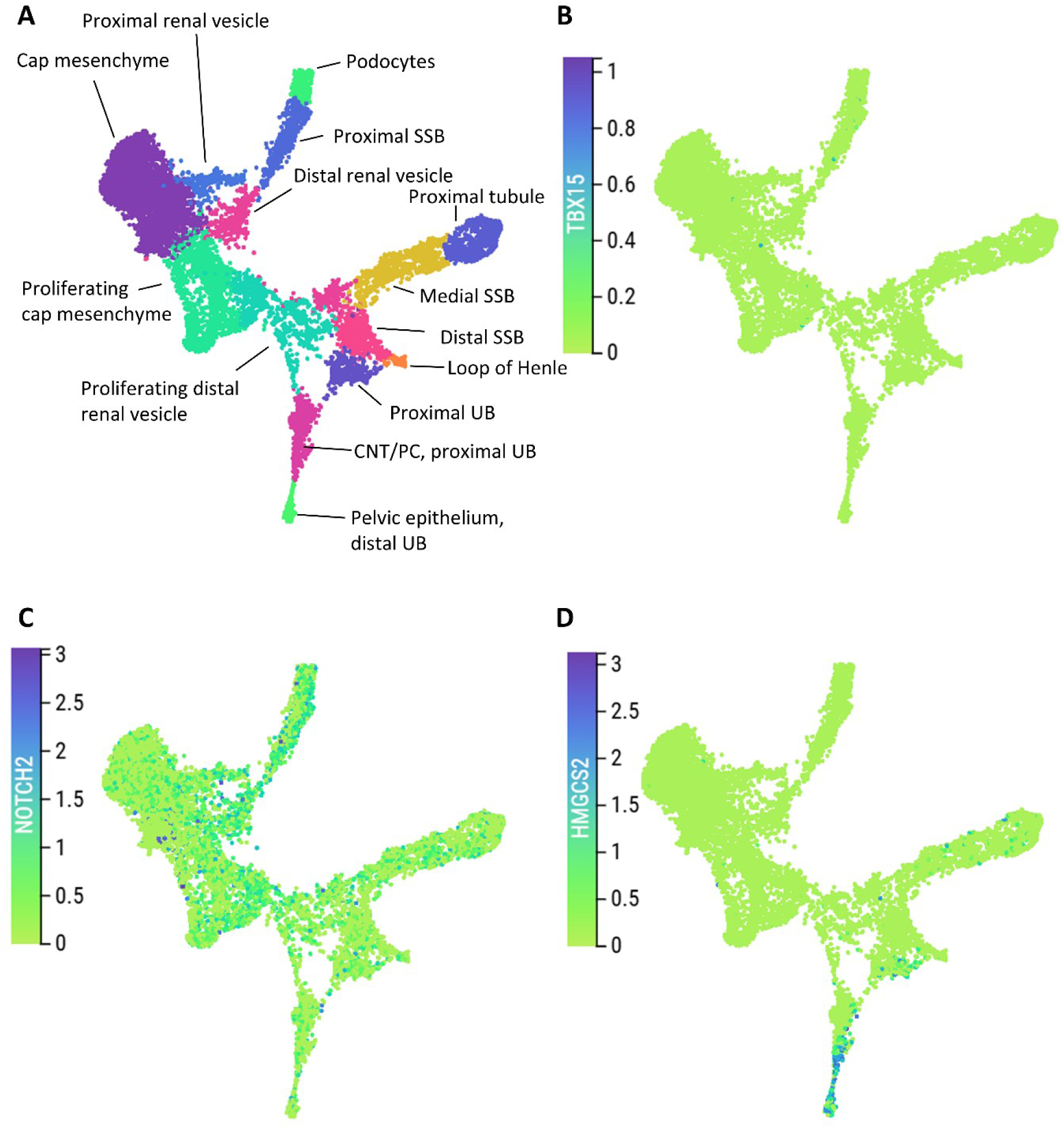
Overview of gene expression of TBX15 (A), NOTCH2 (B) and HMGCS2 (C) in different cell types (D) of the human fetal developing nephron, UB, ureteric bud. SSB, S shaped body. Data source: kidneycellatlas.org/fetal-kidney-developing-nephron

**Supplemental Figure 9.**
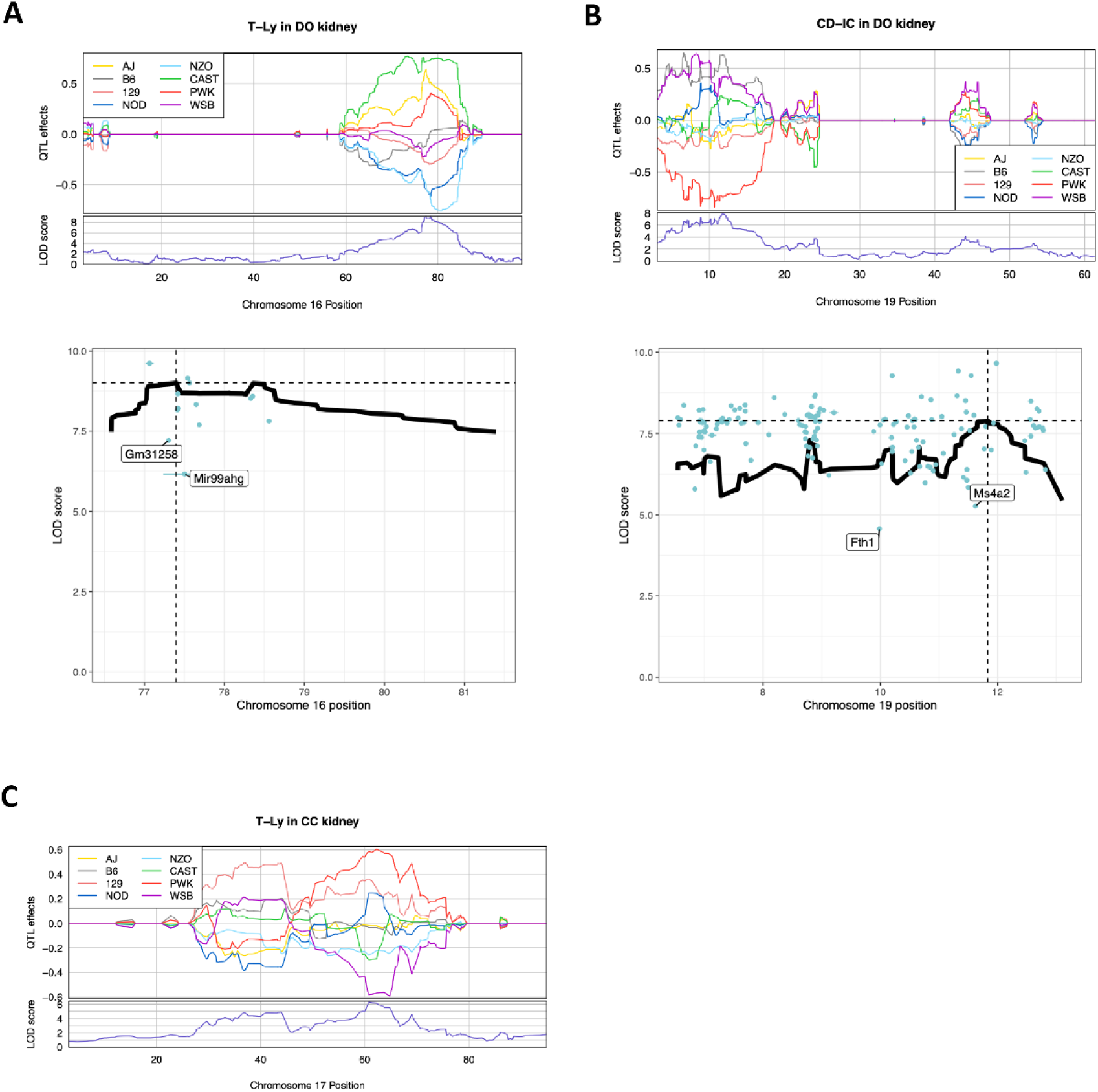
Founder haplotype effects (top) and LOD drop results (bottom) of ccQTL for T-Ly (A) and CD-IC (B), both in DO kidney. For LOD drop plots, thick black line represents the LOD score for the ccQTL support region, peak locus marked by the vertical dashed line and peak LOD score by the horizontal dashed line. Candidate mediator genes represented by blue dot and bar. Candidate mediator genes with notable LOD drops highlighted with labels. (C) Founder haplotype effects of suggestive ccQTL for T-Ly in CC kidney. All ccQTL were highlighted in Figure S5.

